# Genetic architecture influences when and how hybridization contributes to colonization

**DOI:** 10.1101/705954

**Authors:** Bryan Reatini, Todd J. Vision

**Author notes:** Authorship contributions: BR and TJV conceived the work and designed the methodology. BR carried out the simulations and performed data analysis. BR initially drafted, and BR and TJV jointly revised the manuscript.

## Abstract

The role of genetic architecture in adaptation to novel environments has received considerable attention when the source of adaptation variation is *de novo* mutation. Relatively less is known when the source of adaptive variation is inter- or intraspecific hybridization. We model hybridization between divergent source populations and subsequent colonization of an unoccupied novel environment using individual-based simulations in order to understand the influence of genetic architecture on the timing of colonization and the mode of adaptation. We find that two distinct categories of genetic architecture facilitate rapid colonization but that they do so in qualitatively different ways. For few and/or tightly linked loci, the mode of adaptation is via the recovery of adaptive parental genotypes. With many unlinked loci, the mode of adaptation is via the generation of novel hybrid genotypes. The first category results in the shortest colonization lag phases across the widest range of parameter space, but further adaptation is mutation limited. The second category takes longer and is more sensitive to genetic variance and dispersal rate, but can facilitate adaptation to environmental conditions which exceed the tolerance of parental populations. These findings have implications for understanding the origins of biological invasions and the success of hybrid populations.

## Introduction

A persistent question regarding the source of adaptive variation leading to colonization is why intra and interspecific admixture – hereafter collectively referred to as hybridization – sometimes leads to increased colonization success and sometimes does not (Bock et al., 2015; Dlugosch et al., 2015). Hybridization can contribute to colonization by producing an overall increase in standing genetic variation, by transferring adaptive alleles between genetic backgrounds (adaptive introgression), or by producing novel genotypes in hybrids (transgressive segregation) (Pfennig et al., 2016). Many species complexes are known in which hybridization may have contributed to colonization in these ways (Ellstrand and Schierenbeck, 2000; Schierenbeck and Ellstrand, 2008). However, hybridization doesn’t always contribute to increased colonization success, and identifying the genetic factors which may influence when and how hybridization contributes to colonization has become a central objective of the field of invasion genetics (Bock et al., 2015).

A second persistent question in invasion genetics is what the genetic architecture of so-called invasiveness traits tends to be (Bock et al., 2015; Dlugosch et al., 2015). Genetic architecture here is defined as the number of loci governing a quantitative trait, their positions within the genome, and the interactions among different alleles and loci. Much is already known about the genetic architecture of adaptation to heterogeneous environments when adaptation stems from *de novo* mutation. For example, models of adaptation across linear gradients have demonstrated the importance of sufficient genetic variance (Kirkpatrick and Barton, 1997), two patch models have demonstrated the importance of mutations of large effect (Gomulkiewicz et al., 1999; Holt et al., 2003), and a recent evaluation of adaptation across an approximately continuous landscape demonstrated the potential importance of both high genetic variance and mutations of large effect (Gilbert and Whitlock, 2017). However, range expansion may commonly be facilitated by gene flow rather than *de novo* mutation, and studies on the genetic architecture of colonization and range expansion stemming from hybridization are currently lacking.

Here, we wish to examine how genetic architecture itself influences when and how hybridization contributes to colonization success. Hybrid invasions present a unique combination of population genetic forces which may interact with genetic architecture in important ways. For example, selection on beneficial variation within a population receiving migrants may be rendered ineffective due to genetic swamping – the homogenizing effect of persistent gene flow (García-Ramos and Kirkpatrick, 1997; Yeaman and Otto, 2011; Yeaman and Whitlock, 2011). The strength of genetic swamping may be modulated by genetic architecture. Architectures dominated by alleles with large selection coefficients and/or tightly linked loci are more resistant to swamping than those dominated by alleles with small selection coefficients and/or unlinked loci (Tigano and Friesen, 2016; Yeaman and Whitlock, 2011), although architectures characterized by many alleles of small effect may be robust to genetic swamping when genetic variance is sufficiently high (Yeaman, 2015). The implication of this is that, when looking retrospectively at adaptations in systems where there has been persistent gene flow, swamping-resistant genetic architectures may be more likely to be observed.

When hybridization occurs between divergent populations in a new environment, as would be the case between introduced taxa or between an introduced and native taxon, colonization success may depend on the ability to recover and maintain adaptive variation in the face of persistent migration from these divergent source populations. In fact, the time required for natural selection to drive adaptive genetic variation to high frequency is a proposed explanation for one of the most ubiquitous phenomena in invasion biology; the initial lag phase in which introduced populations persist in low numbers before rapid growth (Crooks, 2005; Crooks and Soulé, 1999). Using a source-sink model incorporating adaptation from mutation, Holt et al. (2003) demonstrated that transient occupancy of a sink can persist for a long time before punctuated growth following adaptation.

Given these findings, we hypothesize that genetic architecture, which will influence the recovery of adaptive variation in newly formed hybrid populations, will also influence the duration of the lag phase. If the lag phase is sufficiently long, we may not witness its resolution within our limited window of observation, and conclude that hybridization has failed to contribute to colonization. We hereafter refer to the idea that the genetic architecture of invasiveness traits should influence the contribution of hybridization to colonization – as the *architecture hypothesis of hybrid invasion*.

Here we test the architecture hypothesis of hybrid invasion using individual based simulations. Specifically, we test two underlying predictions: (1) that colonization lag phase duration is sensitive to the genetic architecture of invasiveness traits, (2) that architectures resistant to genetic swamping will produce relatively short lag phases. Given our results, we then test an additional prediction: the filtering effect of colonization on genetic architecture will influence the ability of resulting colonist populations to occupy and adapt to more extreme environments – an important characteristic for hybrid speciation and adaptive radiation.

## Methods

We simulated colonization in a source-sink scenario involving dispersal from two divergent source populations using quantiNemo2 (Neuenschwander et al., 2008). Generations were discrete and consisted of four life cycle events in the following order: reproduction, offspring dispersal, aging, and regulation of adults. We generated polymorphic populations in two separate patches with differing environmental optima. Each initial allele in the population was drawn from a discretized normal distribution with variance specified by the allelic variance of the genetic architecture (Table 1). The populations then experienced a burn-in period with selection and mutation, but no dispersal, in order that they attain mutation-selection-drift equilibrium within each patch. Unless otherwise specified, the burn-in period was 10,000 generations. We then allowed one-way dispersal from these two source populations into a novel environment (the sink) for 500 generations – a timeframe relevant for most contemporary invasions – which we refer to as the *colonization phase*. We assume that occupancy of the novel environment by either source population alone is dependent on recurrent immigration. Thus, it will serve as a sink for maladapted migrants from the source populations, at least initially.

**Table 1.** Genetic architectures tested in simulations of hybrid colonization. Allelic variance is the variance of the distribution of mutation effect sizes. Linkage: unlinked (*U*), one (*W*1) or ten (*W*10) within-trait linkage groups, one (*B*1) or ten (*B*10) between-trait linkage groups, or within-trait and between-trait linkage (*WB*) in which loci from the different traits are interleaved at a distance of 0.5 cM in a single linkage group (see Methods).

| *Loci* | mutation rate | allelic variance | linkage | epistatic variance |
| --- | --- | --- | --- | --- |
| 1 | 10<sup>-4</sup> | 0.5 | *U* | 0 |
| 10 | 10<sup>-4</sup> | 0.05 | *U* | 0 |
| 100 | 10<sup>-4</sup> | 0.005 | *U* | 0 |
| 1000 | 10<sup>-4</sup> | 0.0005 | *U* | 0 |
| 1 | 10<sup>-5</sup> | 0.5 | *U* | 0 |
| 10 | 10<sup>-5</sup> | 0.05 | *U* | 0 |
| 100 | 10<sup>-5</sup> | 0.005 | *U* | 0 |
| 1000 | 10<sup>-5</sup> | 0.0005 | *U* | 0 |
| 10 | 10<sup>-5</sup> | 0.05 | *W1* | 0 |
| 100 | 10<sup>-5</sup> | 0.005 | *W1* | 0 |
| 100 | 10<sup>-5</sup> | 0.005 | *W10* | 0 |
| 1000 | 10<sup>-5</sup> | 0.0005 | *W1* | 0 |
| 1000 | 10<sup>-5</sup> | 0.0005 | *W10* | 0 |
| 10 | 10<sup>-4</sup> | 0.05 | *B1* | 0 |
| 100 | 10<sup>-4</sup> | 0.005 | *B1* | 0 |
| 10 | 10<sup>-4</sup> | 0.05 | *B10* | 0 |
| 100 | 10<sup>-4</sup> | 0.005 | *B10* | 0 |
| 10 | 10<sup>-4</sup> | 0.05 | *WB* | 0 |
| 100 | 10<sup>-4</sup> | 0.005 | *WB* | 0 |
| 10 | 10<sup>-5</sup> | 0.05 | *U* | 0.05 |
| 10 | 10<sup>-5</sup> | 0.05 | *U* | 0.5 |
| 10 | 10<sup>-5</sup> | 0.05 | *U* | 2.25 |
| 100 | 10<sup>-5</sup> | 0.05 | *U* | 0 |

We simulated two quantitative traits and subjected them to stabilizing selection. For the first trait, one source population generated an adaptive genotype during the burn-in period relative to the optimum of the sink patch while the other source population generated a maladaptive genotype (Figure 1). The opposite was true for the second trait, resulting in each source population possessing an adaptive genotype for one trait and a maladaptive genotype for the other with respect to the trait optimum in the sink. In this way, all of the standing genetic variation necessary to adapt to the sink is present in the two source populations at the start of the colonization phase, but it must be assembled via recombination in the sink.

**Figure 1.**
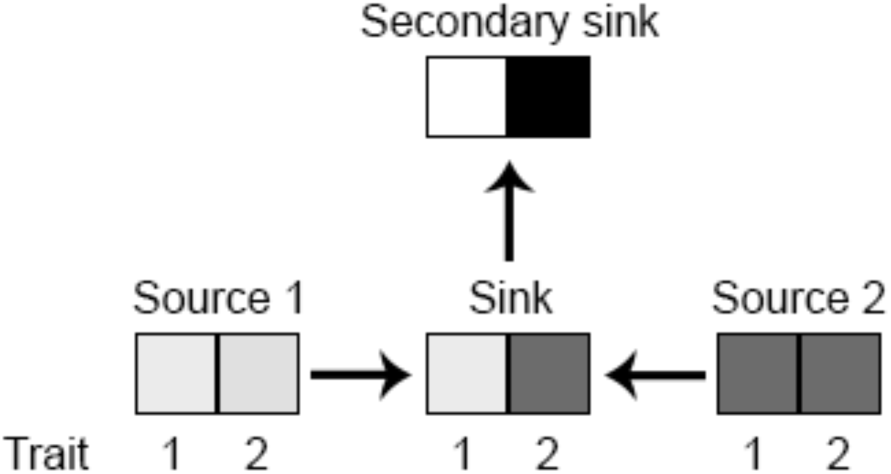
The landscape of adaptive optima for the two quantitative traits. The sink shares optima with one source populations for each trait. The secondary sink has optima more extreme than those of the original source patches.

We evaluated a range of differences in environmental optima (Δ*Z*_opt_ of 6, 8, and 10) between source patches, where *Z*_opt_ represents the optimal phenotype for each quantitative trait. For reference, one unit of *Z*_opt_ is equivalent to two-fold the variance of the mutation effect size distribution for 1-locus architectures. This range is intermediate between smaller values of Δ*Z*_opt_ that yield successful colonization by a single source population in the absence of hybridization, and larger values that fail to colonize under a variety of genetic architectures. Thus, the range of values of Δ*Z*_opt_ we employ encompasses the environmental optima relevant for our analysis of hybridization’s contribution to colonization, given the other assumptions we make about the strength of selection (discussed below).

### Reproduction and dispersal

Reproduction followed a Wright-Fisher Model with a variable population size, *N*, of diploids in which random mating occurred with any individual in the same patch, including self-mating, with probability 1/*N*. Fecundity for each individual was drawn from a Poisson distribution with a mean of four. Growth within each patch was governed by the fecundity of its individuals, and each patch had a carrying capacity of 1000 individuals. We allowed one-way dispersal from source populations into the sink at dispersal rates of *d*=10^−2^ and *d*=10^−3^ during the colonization phase, where dispersal rate is the probability of an offspring migrating to the sink. Our justification for using these rates is as follows; based on our previous assumptions, the total expected number of migrants *N*_m_ entering the sink per generation can be written as *N*_m_ = 2*Kfd*. This simplifies to *N*_m_ = 8000*d*, given that carrying capacity *K* = 1000 for both source populations and fecundity *f* has a mean of four. At *d*=10^−1^, *N*_m_=800, which could result in the sink reaching carrying capacity solely due to the influx and chance survival of migrants, which would not be of interest. At the other extreme of *d*=10^−4^, *N*_m_=0.8, which may not provide sufficient opportunity for hybridization to occur within 500 generation. We therefore limit our analysis to intermediate values of *d*=10^−2^ and *d*=10^−3^, for which *N*_m_=80 and *N*_m_=8, respectively. This provides opportunity for hybridization between migrants while still allowing us to evaluate the effect of an order of magnitude difference in propagule pressure. After dispersal, aging occurred, which removed the adults from each patch and converted the offspring into adults for the next non-overlapping generation. Population regulation then followed, randomly culling the number of new adult individuals in each patch that exceeded the carrying capacity.

### Selection and genetic architecture

Selection acted on offspring survivorship, with an individual’s fitness modeled as its survival probability before aging and reproduction. Stabilizing selection acted on the phenotype of both quantitative traits within each patch. Trait fitness *W*_*t*_ was defined by the standard Gaussian function 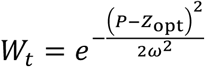, where *P* is the value of the quantitative trait, *Z*_opt_ is the optimal trait value of the patch, and ω^2^ is the width of the fitness function. Following Gilbert et al. (2017), we assume ω^2^ to be 7.5 given a heritability *h*^*2*^ ≈ 1/3 and environmental variance (*V*_*E*_) of one. These values were used throughout all simulations. We then calculate the fitness of an individual *W* as the product of the trait fitness for the two simulated traits: 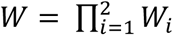.

We simulated quantitative traits under four baseline schemes of genetic architecture characterized by the number of loci (1, 10, 100, and 1000 loci), assuming free recombination, no pleiotropic effects, and additive alleles. We then modified these baseline architectures in subsequent analyses to include variations in linkage and epistasis (Table 1). Mutations acted in a stepwise fashion where the effect of a new mutation was added to the previous allelic value. The variance of the discretized normal distribution from which mutational effects were drawn was scaled by the number of loci underlying the trait (Table 1), such that the effect size of mutations generated during the burn-in period depended upon the model being tested (*i*.*e*., few mutations of large effect for architectures with fewer loci and many mutations of small effect for architectures with many loci). We compared mutation rates of μ=10^−3^, μ=10^−4^, μ=10^−5^, and μ=10^−6^ in order to evaluate the effect of differing values of *V*_*g*_ within source populations on colonization success (Supplemental Figure 1). We then narrowed this range to μ=10^−4^ and μ=10^−5^ for baseline simulations given that these rates bracketed an inflection point in the capacity for adaptation when genetic swamping was strong, consistent with Yeaman (2015). In order to prevent adaptation from mutation during the colonization phase, we only allowed mutation during the burn-in period.

### Analysis

We measured lag phase duration as the number of generations between the initiation of dispersal into the sink (transient occupancy) and the time at which carrying capacity in the sink was reached (permanent residency). We used mean genotypic value, *G*, as a metric to track adaptation within the sink population during the colonization phase. It was calculated as 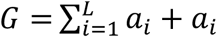 where *L* is the number of loci and *a*_*i*_ and *a*_*i’*_ are the two alleles at the *i*^th^ locus of a diploid individual. This formula is modified in section 3.2.2 to incorporate epistasis.

Given that mutation was not allowed during the colonization phase, sampling the entirety of both source populations at the end of the burn-in period allowed us to determine the origin of every allele in the metapopulation. However, due to the finite number of possible allele effect sizes at each locus (255), there was a small chance that the same allele could be present in both source populations at the end of the burn-in. These identical alleles were ignored. Using these data, we characterized the genetic architecture of adaptation to the sink by measuring the frequency, effect size, and origin of each adaptive allele within the sink immediately after the lag phase. To be included in the analysis, we required an adaptive allele to have a frequency greater than a dispersal-dependent threshold, calculated as the maximum frequency of a new migrant allele entering the sink from one of the two source populations,

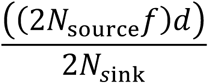

where *f* is fecundity and *d* is dispersal rate. Since *N*_source_ and *N*_sink_ are both *K* after the lag phase and *f* is assumed to have a mean of four throughout all simulations, this simplifies to 4*d*.

We then quantified the relative impact of each of the adaptive alleles in the sink by taking the product of the allele’s frequency and effect size, a quantity we refer to as the *weight* of an allele. By summing the weights of adaptive alleles originating from each source population, we were able to estimate the relative contribution of each source population to adaptation in the sink. When adaptation in the sink proceeds primarily by the recovery of parental genotypes, this is reflected by the total weight of adaptive alleles being primarily from one source. By contrast, when adaptation is achieved through the generation of hybrid genotypes, the contribution from the two source populations is more even. All analyses were averaged across 100 simulation replicates for each scheme of genetic architecture. Analyses of variance (ANOVAs) were then performed to test the effect of the different factors and their interactions on lag time duration. Data and code for reproducing the analyses are available from Dryad (Reatini and Vision, 2020).

## Results

### The ‘when’ and the ‘how’ of hybrid colonization

In order to test the architecture hypothesis of hybrid invasion, we first focused on its primary underlying prediction: colonization lag phase duration is sensitive to the genetic architecture of invasiveness traits. Lag phase duration for the four baseline schemes of genetic architecture (1, 10, 100, or 1000 loci, assuming free recombination and additivity within and between loci) varied with the magnitude of the differences between patch optima (Δ*Z*_opt_), the mutation rate during the burn-in period (which in turn affected the starting values of *V*_*g*_ in the source populations), and the dispersal rate (Figure 2). In mild environments, which we define as having a relatively small Δ*Z*_opt_ of 6, mean colonization lag phase was on the order of ten generations for all genetic architectures and all parameter values, and thus did not differ between architectures. The lag phase when colonizing moderate environments (Δ*Z*_opt_=8) varied depending on the genetic architecture and other assumptions of the model (Figure 3A). In more extreme environments (Δ*Z*_opt_=10), 1-locus architectures were capable of colonizing the sink but all other architectures failed (i.e., the lag phase exceeded 500 generations) for the parameter values we tested. For subsequent analyses, we use Δ*Z*_opt_=8, since for that parameter value, all genetic architectures showed some sensitivity to the other parameters.

**Figure 2.**
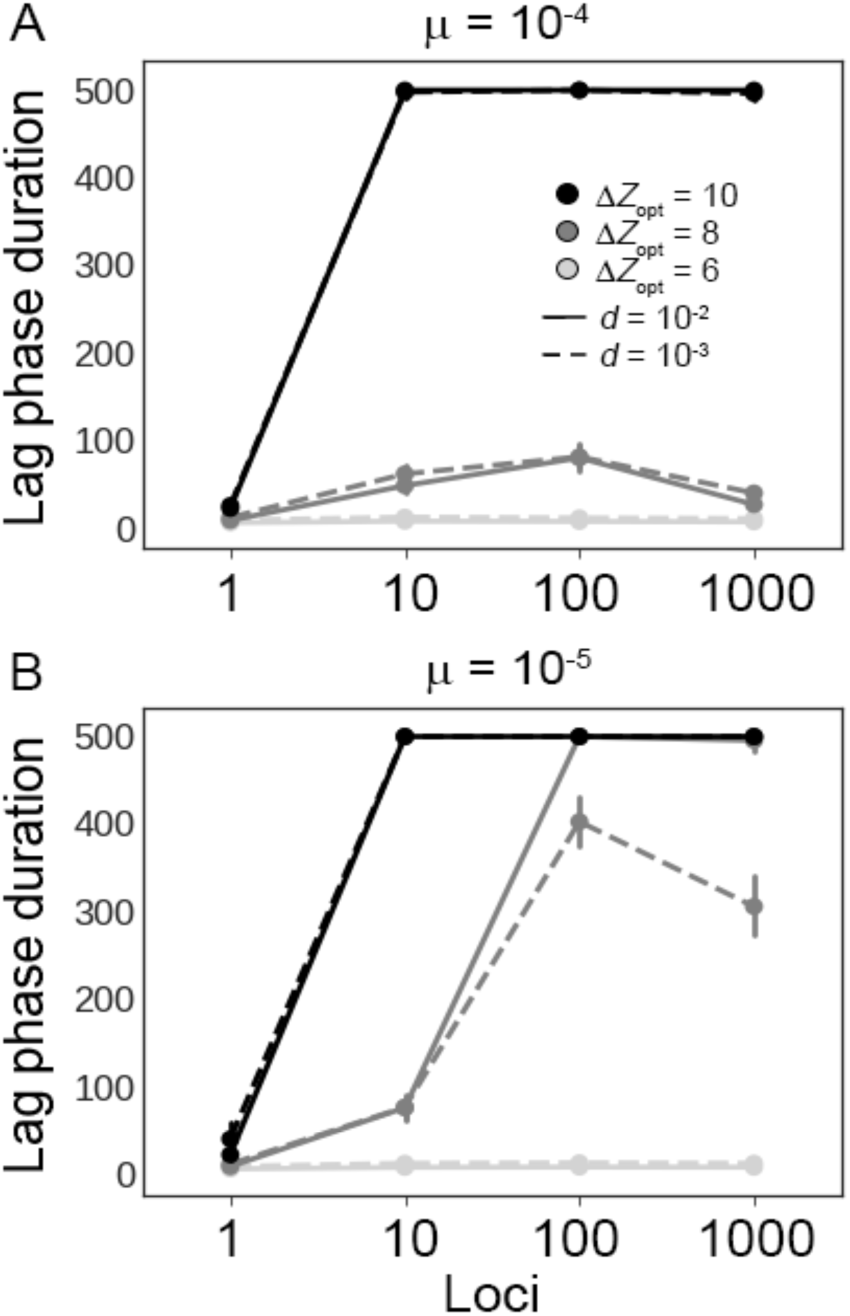
Mean colonization lag phase and 95% confidence intervals for each baseline architecture assuming different values of Δ*Z*_opt_ (light grey=6, dark grey=8, black=10), dispersal rate *d* (solid lines=10^−2^, dashed lines=10^−3^) and mutation rate (μ=10^−4^ in A, μ=10^−5^ in B). For Δ*Z*_opt_=8, significant interactions were found between locus number, mutation rate, and dispersal rate.

**Figure 3.**
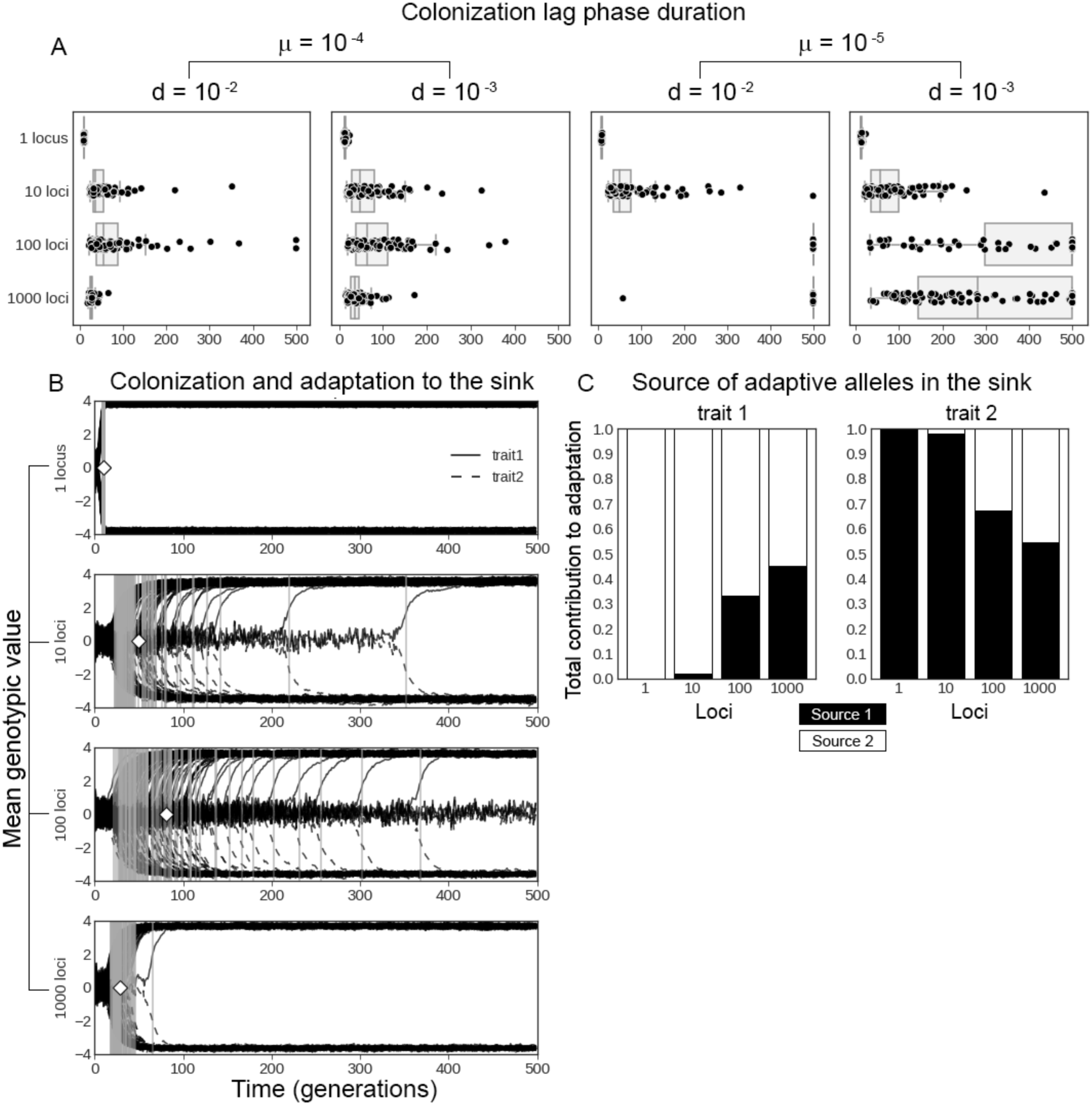
A) Duration of the colonization lag phase (in generations) for each simulation replicate (black dots) under different dispersal rates *d* and mutation rates μ assuming Δ*Z*_opt_=8. B) Recovery of mean genotypic value in the sink for each simulation replicate (black lines), the end of the lag phase for each replicate (grey bars), and mean lag phase duration (white diamonds) – taken from high mutation rate and high dispersal rate simulations from panel A. C) Total contribution to adaptation (in terms of total weight of adaptive alleles) in the sink for each source population averaged across simulation replicates from panel B.

Overall, the lag phase was shorter for the 1-locus simulations compared to all other architectures under all parameter values tested. For architectures with many loci, a low burn-in mutation rate produced longer lag phases and, when combined with high dispersal rate, resulted in a failure to colonize the sink during colonization phase. A significant three-way interaction was found between the number of loci, the dispersal rate, and the mutation rate (Table 2). In order to analyze the underlying factors contributing to this interaction, simple second-order interactions were analyzed by performing two-way ANOVAs between each pair of factors (loci, *μ* and *d*) at each level of the third factor (Table 3). Significant simple second-order interactions were found between mutation and dispersal for 100 and 1000 loci, between loci and dispersal at μ =10^−5^ (Figure 2B), and between loci and mutation at both levels of dispersal. For all architectures, lag phase duration closely corresponded with the timing of adaptation to the sink, as shown for the case of high mutation and dispersal rates in Figure 3B.

**Table 2.** Multifactorial ANOVA for the response variable of colonization lag time and the fixed factors of dispersal rate, mutation rate, and the number of loci.

|  | Type II sum of squares | Degrees of freedom | F | p |
| --- | --- | --- | --- | --- |
| *dispersal* | 39,649 | 1 | 80.495 | < 2.2e-16 |
| *loci* | 17,581,779 | 3 | 1189.814 | < 2.2e-16 |
| *mutation* | 14,335,878 | 1 | 2910.461 | < 2.2e-16 |
| *dispersal-by-loci* | 614,310 | 3 | 41.572 | < 2.2e-16 |
| *dispersal-by-mutation* | 615,323 | 1 | 124.922 | < 2.2e-16 |
| *loci-by-mutation* | 12,812,203 | 3 | 867.042 | < 2.2e-16 |
| *dispersal-by-loci-by-mutation* | 653,133 | 3 | 44.2 | < 2.2e-16 |
| *residual error* | 780,2212 | 1,584 |  |  |

**Table 3.** Two-way ANOVAs for the response variable of colonization lag time and each pair of fixed factors of dispersal rate, mutation rate, and the number of loci at each level of the third factor.

|  | Type II sum of squares | Degrees of freedom | F | p |
| --- | --- | --- | --- | --- |
| dispersal:mutation for 1 locus |  |  |  |  |
| *dispersal* | 951.7 | 1 | 366.129 | < 2.2e-16 |
| *mutation* | 1.8 | 1 | 0.701 | 0.403 |
| *dispersal:mutation* | 5.1 | 1 | 1.948 | 0.164 |
| *residuals* | 1029.4 | 396 |  |  |
| dispersal:mutation for 10 loci |  |  |  |  |
| *dispersal* | 4823 | 1 | 1.418 | 0.23448 |
| *mutation* | 44711 | 1 | 13.143 | 0.00033 |
| *dispersal:mutation* | 4090 | 1 | 1.202 | 0.27356 |
| *residuals* | 1347178 | 396 |  |  |
| dispersal:mutation for 100 loci |  |  |  |  |
| *dispersal* | 229393 | 1 | 26.52 | 4.12e-07 |
| *mutation* | 13671876 | 1 | 1580.65 | < 2.2e-16 |
| *dispersal:mutation* | 241228 | 1 | 27.89 | 2.12e-07 |
| *residuals* | 3425205 | 396 |  |  |
| dispersal:mutation for 1000 loci |  |  |  |  |
| *dispersal* | 775632 | 1 | 101.4 | < 2.2e-16 |
| *mutation* | 13431492 | 1 | 1756.1 | < 2.2e-16 |
| *dispersal:mutation* | 1023132 | 1 | 133.8 | < 2.2e-16 |
| *residuals* | 3028799 | 396 |  |  |
| loci:dispersal at $\mu=10^{-4}$ | | | | |
| *loci* | 539545 | 3 | 91.015 | < 2.2e-16 |
| *dispersal* | 11974 | 1 | 6.06 | 0.014 |
| *loci:dispersal* | 6101 | 3 | 1.029 | 0.379 |
| *residuals* | 1565009 | 792 |  |  |
| loci:dispersal at $\mu=10^{-5}$ | | | | |
| *loci* | 29854436 | 3 | 1263.64 | < 2.2e-16 |
| *dispersal* | 999839 | 1 | 126.96 | < 2.2e-16 |
| *loci:dispersal* | 1261342 | 3 | 53.39 | < 2.2e-16 |
| *residuals* | 6237202 | 792 |  |  |
| loci:mutation at $d=10^{-2}$ | | | | |
| *loci* | 11845016 | 3 | 1966 | < 2.2e-16 |
| *mutation* | 10445649 | 1 | 5202 | < 2.2e-16 |
| *loci:mutation* | 9299236 | 3 | 1544 | < 2.2e-16 |
| *residuals* | 1590229 | 792 |  |  |
| loci:mutation at $d=10^{-3}$ | | | | |
| *loci* | 6351073 | 3 | 269.9 | < 2.2e-16 |
| *mutation* | 4505552 | 1 | 574.4 | < 2.2e-16 |
| *loci:mutation* | 4166099 | 3 | 177.1 | < 2.2e-16 |
| *residuals* | 6211983 | 792 |  |  |

In addition to quantitatively affecting the duration of the lag phase, the different genetic architectures led to qualitatively different modes of adaptation to the sink (Figure 3C). We analyzed the genetic architecture of adaptation under high mutation and dispersal rates, since the lag phase duration was minimized with these parameter values for all genetic architectures. For the 1-locus and 10-locus simulations, adaptation to the sink was based on the same alleles that contributed to adaptation in the source populations. Specifically, 100% of alleles contributing to adaptation in the sink originated from their respective adapted source population for 1-locus architectures. For 10-locus simulations, the mean contribution across replicates was 98.2% for both traits. In contrast, adaptation to the sink for 100-loci and 1000-loci architectures proceeded predominantly by the generation of novel hybrid genotypes. For 100-loci, the mean contribution of adaptive alleles originating from source population one was 33.1% for trait one and 67.1% for trait two. For 1000-loci, the mean contribution from source population one was 45.3% for trait one and 54.5% for trait two. Thus, two modes of adaptation in the sink are observed; the recovery of parental genotypes as seen for 1 and 10-locus architectures, and the generation of hybrid genotypes for 100 and 1000-loci architectures.

### Linkage

Above, we have assumed free recombination between all loci and additive contributions to phenotype for the sake of simplicity, but both linkage and epistasis may influence adaptation in newly formed hybrid populations (Abbott et al., 2013; Harrison and Larson, 2014; Simon et al., 2018; Yeaman and Whitlock, 2011). Linkage between loci reduces the recombination rate, and therefore may reduce the strength of genetic swamping when there are many loci (Tigano and Friesen, 2016). Since genetic swamping from source populations appears to be the force that impeded colonization for 100-loci and 1000-loci architectures when the burn-in mutation rate was low, we hypothesized that linkage among the loci within each adaptive trait could rescue these architectures under such conditions. To test this hypothesis, we incorporated varying degrees of within-trait linkage into 10, 100, and 1000 locus architectures (Table 1). All loci for each trait were clustered within either one or ten linkage groups 100 cM in length. Loci were spaced at 1 cM intervals in all cases except that of 1000 loci on a single linkage group, in which spacing was 0.1 cM. Importantly, there was free recombination between the loci of the two different traits in these simulations.

In order to evaluate the effect of selective interference between traits on lag phase duration, we then simulated linkage between the loci of the two different traits for 10 and 100-locus architectures. First, we tested the effects of between-trait linkage alone by placing pairs of loci, one for each trait, at a distance of 1 cM or 10 cM away from one another on each chromosome. Thus, there were 10 and 100 chromosomes for 10 and 100-locus architectures, respectively. In this way, we simulated linkage between the loci of the two different traits while preventing linkage among loci of the same trait. We then evaluated the combined effects of within-trait linkage and between-trait linkage by simulating 10 and 100-locus architectures in which all loci for both traits were interleaved at 0.5 cM intervals on a single 100 cM chromosome. These simulations were run under conditions of a low burn-in mutation rate (*μ*=10^−5^) and high rate of dispersal (*d*=10^−2^).

Incorporating within-trait linkage rescued the 100-loci and 1000-loci architectures, with one locag group resulting in shorter lag phases than ten (Figure 4A). Within-trait linkage also resulted in a shift in the mode of adaptation away from the generation of novel hybrid genotypes and towards the recovery of parental genotypes (Figure 4B). Between-trait linkage alone (ie. without within-trait linkage) resulted in 100% of replicates failing to colonize the sink for 10-locus architectures. For 100-locus architectures, the percentage of replicates which failed to colonize the sink increased from 2% in baseline simulations to 9% and 14% for 1 cM and 10 cM simulations, respectively. For interleaved architectures, with both between-trait and within-trait linkage, 98% and 80% of replicates failed to colonize the sink for 10-locus and 100-locus architectures, respectively.

**Figure 4.**
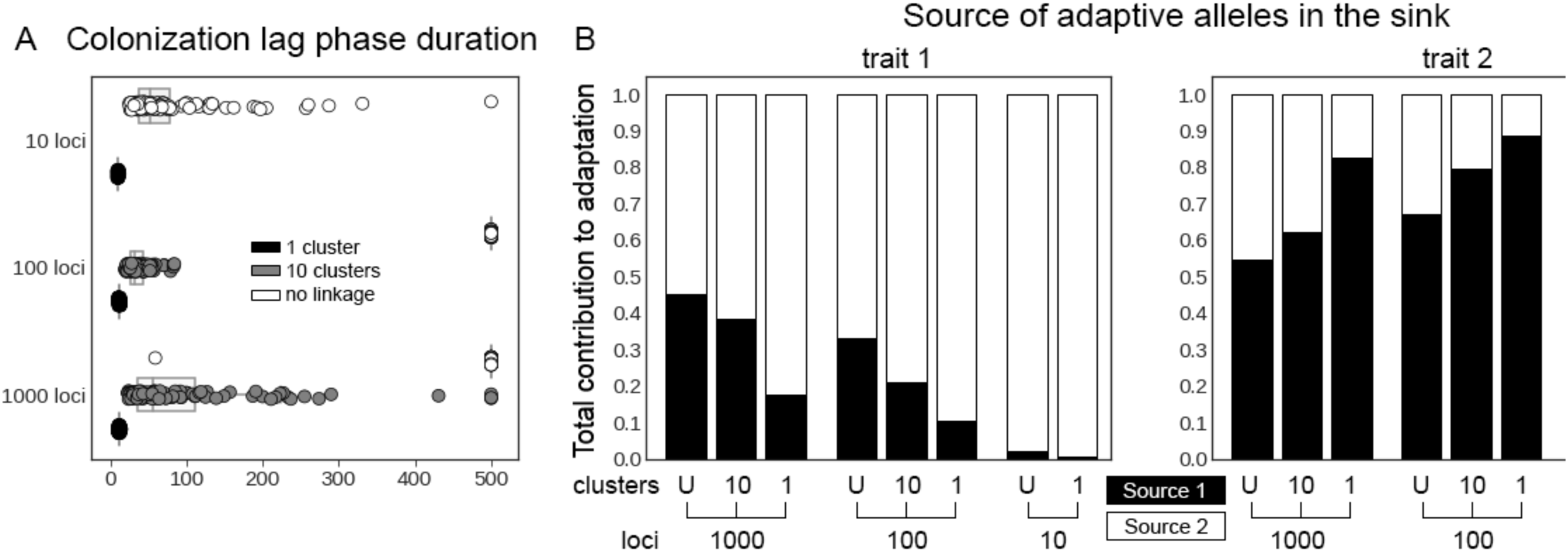
A) Duration of colonization lag phase for each simulation replicate incorporating varying degrees of linkage and assuming low burn-in mutation rate (μ=10^−5^) and high dispersal (*d*=10^−2^). B) Genetic architecture of adaptation to the sink assuming 1 or 10 linkage groups from panel A in comparison to baseline simulations with no linkage (U=unlinked) assuming high mutation rate (μ=10^−4^), and high dispersal rate (*d*=10^−2^).

### Epistasis

Epistasis may influence adaptation in a hybrid zone by creating novel interactions between divergent migrant genotypes. If epistatic interactions are favored by selection, epistasis may result in shorter lag phases and a shift in the mode of adaptation to the generation of novel genotypes. While if such interactions are disfavored, lag phase duration may be lengthened while parental genotypes are recovered. In order to test these predictions, we incorporated epistasis into the model such that the genotype of an individual was determined by the combination of the additive effects of all alleles and an epistatic effect unique to each multilocus genotype. The genotypic value of an individual *G* was computed as 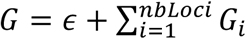 where *ϵ* represents the epistatic effect of the individuals genotype, and *G*_*i*_ represents the additive effect of locus *i*. The epistatic effect for each unique multilocus genotype was drawn from a discretized normal distribution, and we tested a range of variances of that distribution (*V*_*ϵ*_=0.05, 0.5, 2.25). The lowest value in this range reflects the variance of the distribution from which mutation effects were drawn for baseline 10-locus simulations, the middle value represented that base variance multiplied by the ten loci, and the highest value represented that base variance multiplied by the number of pairwise combinations between ten loci. We constrained our test of epistasis to ten locus simulations because computing the epistatic effects for 100- and 1000-loci simulations was not computationally feasible. For the same reason, we also limited the maximum number of alleles at each locus to 55, the minimum value possible within quantiNemo2. We assumed the lower burn-in mutation rate and the higher dispersal rate, since these were the conditions for which there was strong genetic swamping in previous simulations.

Overall, the inclusion of epistasis resulted in a greater percentage of replicates that failed to colonize the sink: 11%, 33%, and 19% of replicates for *V*_*ϵ*_=0.05, 0.5, 2.25, respectively, in comparison to 1% for the baseline architecture without epistasis. Incorporating epistasis resulted in significantly greater variance in colonization lag phase (σ^2^=2.3e4, 4.3e4, and 3.4e4) for *V*_*ϵ*_=0.05, 0.5, and 2.25 respectively) compared to the baseline architecture (σ^2^=5.4e3) (Table 4, Figure 5A). Given the heterogeneity of variance, Games-Howell *post hoc* tests were performed to compare the means. Lag phase duration did not differ significantly among the nonzero values of *V*_*ϵ*_, but mean lag phase duration was significantly longer for the two highest values of *V*_*ϵ*_=0.5 and *V*_*ϵ*_=2.25 in comparison to the baseline architecture without epistasis (Table 4). We did not observe a shift in the mode of adaptation for those replicates that did succeed in colonizing the sink (Figure 5B).

**Table 4.** Effects of epistasis on colonization lag time. Top: Levene’s test for homogeneity of variance for varying levels of epistasis (*V*_*ϵ*_ = 0, 0.05, 0.5, and 2.25). Bottom: pairwise comparisons of means using *post hoc* Games-Howell tests.

|  | Degrees of<br>freedom | F | p |  |  |  |
| --- | --- | --- | --- | --- | --- | --- |
| epistasis | 3 | 8.671 | 1.39e-05 |  |  |  |
| residual error | 396 |  |  |  |  |  |
|  | Post hoc Games-Howell tests |  |  |  |  |  |
| pair | difference | lower | upper | t | degrees of<br>freedom | p |
| 0.05-0 | 31.66 | -12.45 | 75.77 | 1.87 | 142.13 | 0.247 |
| 0.5-0 | 96.87 | 39.64 | 154.10 | 4.41 | 123.40 | 1.3e-4 |
| 2.25-0 | 53.49 | 1.79 | 105.19 | 2.69 | 129.44 | 3.96e-2 |
| 0.5-0.05 | 65.21 | -1.57 | 131.99 | 2.53 | 182.26 | 5.84e-2 |
| 2.25-0.05 | 21.83 | -40.32 | 83.98 | 0.91 | 191.41 | 0.799 |
| 2.25-0.5 | -43.38 | -115.28 | 28.52 | 1.56 | 195.43 | 0.402 |

**Figure 5.**
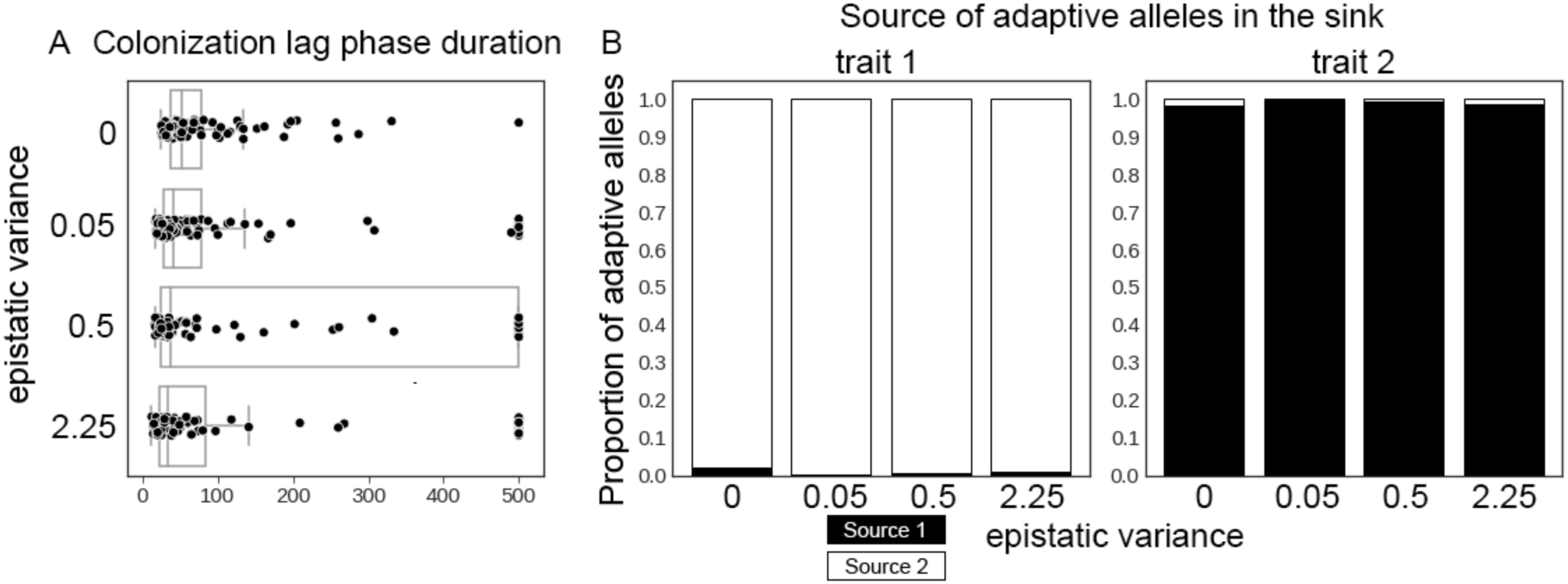
A) Duration of colonization lag phase for each simulation replicate for the three values of epistatic variance (*V*_*ϵ*_) compared with the baseline simulations with no epistasis and assuming a low burn-in mutation rate (μ=10^−5^) and high dispersal (*d*=10^−2^). B) Proportion of adaptive alleles originating from each source population for the three values of *V*_*ϵ*_ in comparison to baseline simulations with no epistasis.

### Introgression vs. transgressive segregation

So far, we have interrogated how genetic architecture may influence the timing of colonization and mode of adaptation by hybrid populations, but we have not considered post-colonization range expansion and evolution. In section 3.1, we observed two primary, and mutually exclusive, modes of adaptation to the sink: either the recovery of parental genotypes within a trait (as seen for 1- and 10-locus architectures) or the generation of hybrid genotypes (as seen for 100- and 1000-locus architectures). Since we assumed additive contributions to the genotypic value for those baseline simulations, we can infer that complementary gene action explains the presence of adaptive alleles from both sources for 100- and 1000-locus architectures. We use complementary gene action here in the sense of alleles from different parents that share the same direction of effect being combined into a multilocus genotype (Rieseberg et al., 1999). This leads us to predict that, for 100 and 1000-locus architectures, but not for 1 and 10-locus architectures, hybrid populations will be capable of further adaptation to optima exceeding that of either source patch.

In order to test this prediction, we conducted simulations in which an additional patch (a secondary sink) was added adjacent to the primary sink with an environmental optimum twice that of the primary sink for each trait (Figure 1, Δ*Z*_opt_=16 for the secondary sink). We allowed one-way dispersal from the primary to the secondary sink. We then monitored colonization and adaptation to the secondary sink by tracking occupancy and mean genotypic value over time. Since mutation was prohibited during the colonization phase, a hybrid population capable of adapting to the extreme optima of the secondary sink would necessarily do so via transgressive segregation of standing variation. We tested baseline architectures in the absence of linkage assuming a high burn-in mutation rate (μ=10^−4^) and high dispersal rate (*d*=10^−2^), since these parameters yielded the shortest lag phases across all architectures in section 3.1.

In agreement with our prediction, we found that the 100-locus and 1000-locus architectures were capable of colonizing the secondary sink while the 1-locus and 10-locus architectures were unable to do so, and that colonization of the secondary sink corresponded with adaptation to its extreme optima (Figure 6).

**Figure 6.**
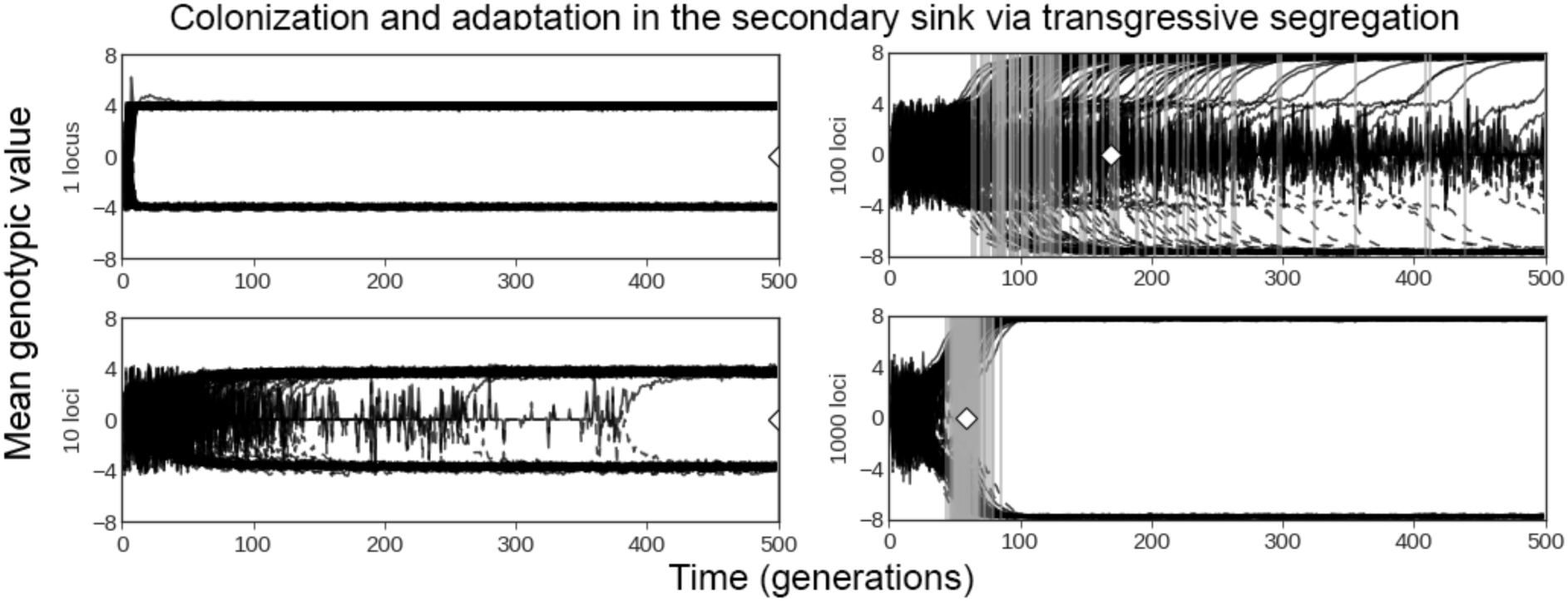
Mean genotypic value in the secondary sink plotted over time for each simulation replicate (black lines). Genotypic values of 4/-4 represent the optima of source patches, whereas genotypic values of 8/-8 represent the optima of the secondary sink. Grey bars denote when carrying capacity was reached in the secondary sink for each simulation replicate, and white diamonds mark the mean.

The amount of transgressive segregation due to complementary gene action may relate to divergence, with increasing complementation resulting from the accumulation of fixed differences between populations over time (Bock et al., 2015; Rieseberg et al., 1999). Increased divergence may therefore reduce the colonization lag phase for architectures that rely on transgressive segregation (*e*.*g*., 100-locus architectures), whereas it may not influence the lag phase for architectures that rely on adaptive introgression (*e*.*g*., 10-locus architectures). We tested the effect of divergence on colonization lag phase by extending the burn-in period from 10,000 generations to 20,000 or 50,000 generations for baseline 10 and 100-locus architectures with a low burn-in mutation rate (μ=10^−5^) and a low dispersal rate (*d*=10^−3^). As predicted, increasing the burn-in period resulted in a slight increase in complementation and a reduction in the mean lag phase duration for 100-locus architectures, but had little effect on 10-locus architectures (Figure 7).

**Figure 7.**
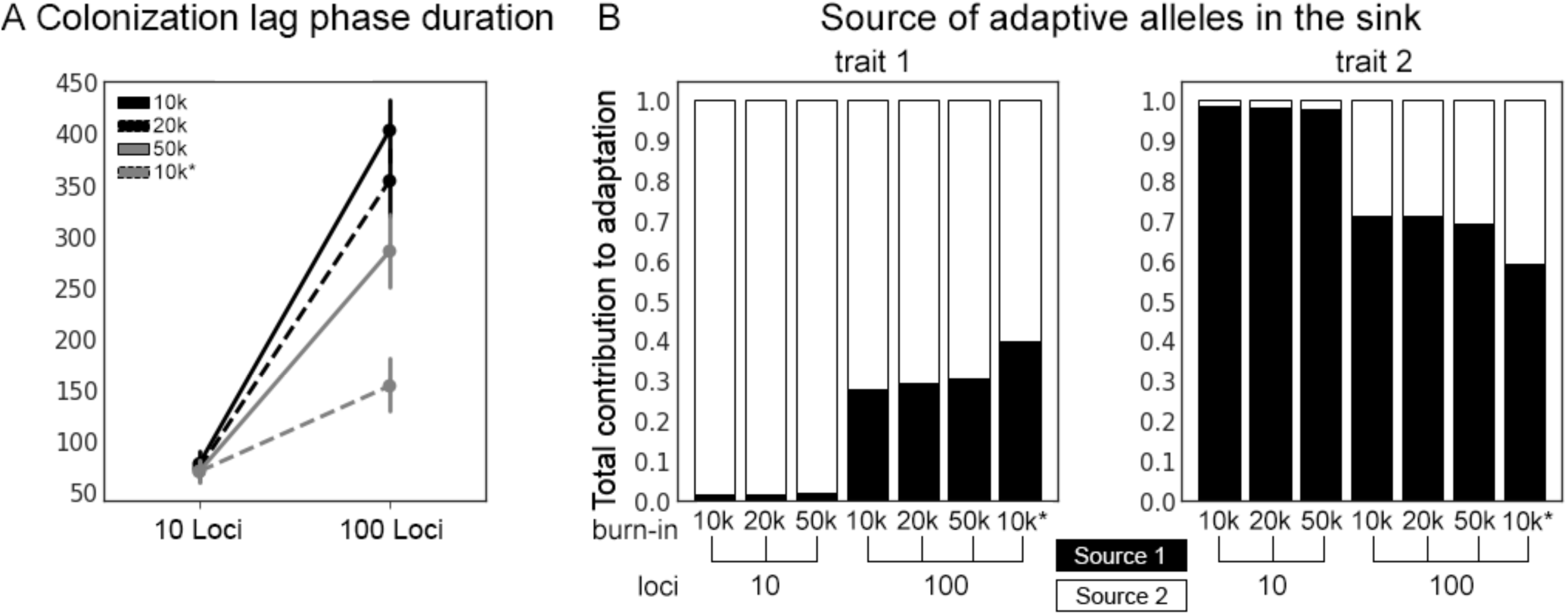
A) Colonization lag phase duration for 10-locus and 100-locus architectures for burn-in durations of 10k generations, 20k generations, and 50k generations, and for allelic variance = 0.05 (10k*). B) Source of adaptive alleles contributing to adaptation in the sink for the same simulations as in A.

Given that large effect alleles are predicted to be more resistant to genetic swamping than small effect alleles (Tigano and Friesen, 2016; Yeaman and Whitlock, 2011), the consequences of increasing the burn-in time might be due, at least in part, to allowing time for larger-effect mutations to arise. In order to evaluate the role of effect size on its own, we increased the variance of the mutational effect sizes (allelic variance) for 100-locus architectures to match that of baseline 10-locus architectures (Table 1). The larger effect size resulted in increased complementation and a corresponding reduction in the mean lag phase duration for 100-locus architectures (Figure 7). However, lag phase duration was still shorter for 10-locus architectures with the same allelic variance (Figure 7A).

## Discussion

The steady accumulation of examples of colonization and invasion by hybrid populations in recent decades (Ellstrand and Schierenbeck, 2000; Schierenbeck and Ellstrand, 2008) has made it a priority in the field of invasion genetics to understand the genetic factors underpinning the contribution of hybridization to colonization (Bock et al., 2015). It is now understood that hybridization can contribute to colonization by producing an overall increase in standing genetic variation, the transfer of adaptive alleles between genetic backgrounds (adaptive introgression), and/or the production of novel genotypes in hybrids (transgressive segregation) (Pfennig et al., 2016).

Our aim here was to test, using individual based simulations, how the architecture of invasiveness traits can influence the contribution of hybridization to colonization. We focused on two underlying predictions: (1) that colonization lag phase duration would be sensitive to genetic architecture and (2) that architectures resistant to genetic swamping would produce shorter lag phases. Given our results, we then tested an additional prediction that the filtering effect of colonization on genetic architecture would influence the ability of resulting colonist populations to occupy and adapt to more extreme environments.

We found two distinct categories of genetic architecture that facilitate rapid colonization by hybrid populations, doing so in fundamentally different ways. The first category included architectures with few loci and/or tightly linked loci. The mode of adaptation observed for such architectures was the recovery of adaptive parental genotypes. Because this pattern is akin to adaptive introgression, we refer to this as an ‘adaptive introgression-type architecture’. The second category included architectures with many freely recombining loci (no within-trait linkage or between-trait linkage) and high *V*_*g*_, in which the mode of adaptation was the generation of novel hybrid genotypes. Because this second pattern is more akin to transgressive segregation, we refer to this as a ‘transgressive segregation-type architecture’.

We found that the extremes within these categories produced the shortest lag phases, with a single locus or a single linkage group producing shorter lag phases than 10 loci or 10 linkage groups for ‘adaptive introgression’-type architectures, and 1000 loci producing shorter lag phases than 100 loci for ‘transgressive segregation’-type architectures. We conclude that lag phase duration is predominantly driven by the number of loci underlying a trait, modulated by their position within the genome and effect size of alleles. These results support our first prediction that lag phase duration is sensitive to the genetic architecture of invasiveness traits.

The second prediction, that genetic architectures resistant to genetic swamping produce shorter lag phases, is supported by two lines of evidence. First, when *V*_*g*_ in the source populations was high (burn-in μ=10^−4^), architectures with many freely recombining loci succeeded in colonization at both low and high dispersal rates, but when *V*_*g*_ in source populations was low (burn-in μ=10^−5^), increased dispersal resulted in colonization failure across all replicates. The increase in lag duration with increased dispersal suggests that the establishment of adaptive genotypes was hindered by a greater influx of maladaptive alleles. This is consistent with previous studies of adaptation across clines that have shown that such architectures are vulnerable to genetic swamping (Yeaman and Otto, 2011; Yeaman and Whitlock, 2011). Our observation that high *V*_*g*_ counteracted increased dispersal, and in fact reduced lag phase significantly for 1000-locus architectures, is consistent with Yeaman (2015), who found that the effects of genetic swamping may be mitigated by sufficiently high *V*_*g*_ for architectures dominated by alleles of small effect. Our model demonstrates that this mitigating effect of *V*_*g*_ on the strength of genetic swamping can significantly reduce lag phase duration for hybrid invasions with such architectures.

The second line of evidence supporting the prediction that genetic architectures resistant to genetic swamping produce shorter lag phases is that incorporating within-trait linkage rescued some simulations in which colonization would otherwise fail. In particular, for the many-locus architectures, when *V*_*g*_ was low and dispersal was high, colonization failed in the case of free recombination but succeeded in the presence of within-trait linkage as long as between-trait linkage was absent. Incorporating within-trait linkage also resulted in a corresponding shift in the mode of adaptation towards the recovery of parental genotypes. This suggests that by reducing recombination between loci, linkage facilitates the recovery of adaptive parental haplotypes despite a steady influx of maladaptive migrant alleles, which is consistent with previous research demonstrating that linkage mitigates the effects of genetic swamping in hybridizing populations (Yeaman and Whitlock, 2011).

A consequence of the longer lag phases observed for architectures prone to swamping is that colonization of the sink often failed to occur within our limited window of observation. For instance, many-locus architectures failed to colonize the sink when *V*_*g*_ was low and dispersal was high. This suggests that these architectures may be under-represented among those contributing to successful hybrid invasions, particularly for recent introductions and organisms with long generation times, because the time since introduction would often be too short. Our results also suggest that, for those hybrid invasions that *are* successful, invasion-related traits would be enriched for ‘adaptive introgression’-type architectures, since these facilitate rapid colonization across the widest range of parameter space.

Strong epistatic interactions may, on average, hinder colonization despite accelerating it in some instances. For 10-locus architectures, we found that the inclusion of epistasis produced greater variance in colonization lag phase and a greater percentage of replicates that failed to colonize the sink. Lag phase was significantly longer when epistatic variance was high in comparison to baseline architectures without epistasis. Interestingly, epistasis had no observable effect on the mode of adaptation to the sink. Further study will be needed to determine if these findings can be generalized to architectures with more loci.

Our results point to an interesting a tradeoff between the modes of adaptation associated with the two extremes of swamping-resistant architectures. ‘Adaptive introgression’-type architectures produce the shortest invasion lag phases across the widest range of parameter space, with 1-locus and single linkage group architectures producing lags with mean duration on the order of ten generations (Figures 2 and 4). Such a capacity to rapidly colonize environments within the bounds of parental trait values may result in initial invasions that spread rapidly. However, such architectures necessarily limit adaptive traits to those that can be obtained by reconstituting parental genotypes, and further adaptation to novel environments beyond the bounds of parental phenotypes may be mutation-limited (Figure 6). By contrast, ‘transgressive segregation’-type architectures produce relatively longer lag phases and fail to colonize the sink under some conditions of *V*_g_ and dispersal. Once established, however, these hybrid populations demonstrate a capacity to adapt to environmental conditions beyond the range of parental populations. Transgressive segregation-type architectures thus may facilitate invasions that can spread into a wider variety of novel habitats (Figure 6).

Another difference between the genetic architectures is in their sensitivity to linkage between traits. In general, we found that selective interference between traits hindered colonization, but that its effect was more pronounced for adaptive introgression-type architectures than transgressive segregation-type architectures.

As noted recently by Marques *et al*. (2019), admixture-derived large-effect haplotypes and the generation of novel genotypes via transgressive segregation are may be frequent sources of adaptive variation for speciation and adaptive radiation. While our simulations do not interrogate the contribution of admixture variation to reproductive isolation *per se*, the colonization of novel or extreme environments by hybrids can promote ecological differentiation between hybrid and parental populations, which in turn fosters conditions favorable for speciation and adaptive radiation (Abbott et al., 2013; Seehausen, 2004). Given our finding that only successful hybrid colonists with ‘transgressive segregation’-type architectures were capable of further colonization and adaptation to more extreme environments, we would predict that such colonists are more likely to generate new hybrid species and adaptive radiations than those with ‘adaptive introgression’-type architectures.

We recognize that the model we present is a crude representation of the complexity of a hybrid invasion, and we have made many simplifying assumptions in order to focus on the role of genetic architecture. For one, we have simulated two source populations with identical population sizes and dispersal pressure into the sink. The demographic histories and extent of gene flow will generally differ between hybridizing populations. The symmetric hybridization scheme such as the one we have explored would be expected to produce the shortest possible lag times given that variation from both source populations is required for adaptation in both traits and that the two traits are of equal importance to fitness. Additionally, hybridization may shift from being beneficial at the establishment phase to maladaptive when persistent migration occurs between core and peripheral populations (García-Ramos and Kirkpatrick, 1997). We explore the effect of genetic swamping during the establishment phase alone, but it is likely to be a persistent force which shapes the adaptive trait architecture of hybrid colonists throughout invasion.

Second, we assume the two source populations are recently diverged. Divergence may influence the contribution of hybridization to colonization in at least two ways. As observed for transgressive segregation-type architectures, increased divergence may increase complementary gene action and provide the allelic variation necessary to generate successful novel genotypes in hybrids. Thus, at least for the relatively short divergence times we tested, colonization success appears to increase with increased divergence for transgressive segregation-type architectures. However, increased divergence may also lead to the accumulation of genetic incompatibilities (ie. Dobzhansky-Muller Incompatibilities or DMIs) between the source populations. DMIs are known to influence the genomic landscape of introgression (Comeault, 2018; Harrison and Larson, 2014), and undoubtedly play an important role in determining when and how hybridization contributes to colonization success. We do not include the evolution of DMIs in our model, and therefore we chose to limit our burn-in period to a timeframe relevant to recently diverged taxa in which DMIs would be less likely to be observed. Models involving longer periods of isolation and the evolution of DMIs will be necessary to comprehensively evaluate the influence of divergence on the contribution of hybridization to colonization.

A third assumption is that we can consider the two traits of interest in isolation from other traits not directly related to colonization. Empirical evidence does suggest that range expansions may commonly involve relatively few adaptive traits (Bock et al., 2015), but it may be important to consider both interference between those traits – as we have done – and, especially in hybrids, other traits under selection in the hybrids. With many segregating polymorphisms under selection, there is considerable opportunity for selective interference (Comeron and Kreitman, 2002; McVean and Charlesworth, 2000), which could delay or prevent adaptation to a novel environment. In a hybrid, segregating loci with formerly fixed differences, affecting traits unrelated to colonization *per se*, may also be newly visible to selection. Thus, not only does selective interference between loci from the two different traits potentially hinder colonization, but this would be exacerbated in a more realistic scenario with many more traits segregating for genetic variation in the hybrids.

A fourth assumption is that new mutations do not accumulate during the colonization phase and thus do not contribute to colonization of the sink. Range expansion stemming from *de novo* mutation has been thoroughly studied theoretically (Gilbert and Whitlock, 2017; Gomulkiewicz et al., 1999; Holt et al., 2003; Kirkpatrick and Barton, 1997), and could feasibly contribute to adaptation within novel environments on timescales relevant for contemporary invasions (Bock et al., 2015; Dlugosch et al., 2015). Moreover, the time required for beneficial mutations to accumulate is another potential genetic cause of invasion lag phases (Crooks and Soulé, 1999; Holt et al., 2003). In reality, hybrid invasions may involve adaptations arising from a combination of mutation and gene flow, and our model only focuses on the latter.

A final assumption is that the lag phase is limited only by local adaptation. Although this was a necessary assumption in order to interrogate the potential role of genetic architecture on lag phase duration, there are other factors that could contribute to the invasion lag phase. These include overcoming Allee effects (Aikio et al., 2010), the time to reach sexual maturity (Wangen and Webster, 2006), changes to the receiving environment (Crooks, 2005; Crooks and Soulé, 1999), and, as discussed above, the accumulation of new mutations.

Despite these simplifying assumptions, our results allow us to make a number of robust conclusions. Primarily, we demonstrate that genetic architecture can influence when and how hybridization contributes to colonization. We find that two categories of swamping-resistant genetic architectures produce short colonization lag phases, and that they do so by qualitatively different modes of adaptation. We also find that there is a tradeoff between the two categories of colonist genetic architectures which may influence future adaptation to more extreme environments.

## Acknowledgements

We thank Morgan Holder and Wesley Price for assistance in conducting initial explorations of the simulation model and software.

## Data and software availability

Simulation settings files and custom analysis code will be made available at the Dryad Digital Repository (https://doi.org/10.5061/dryad.3xsj3txbz).

---

**Supplemental Figure 1.**
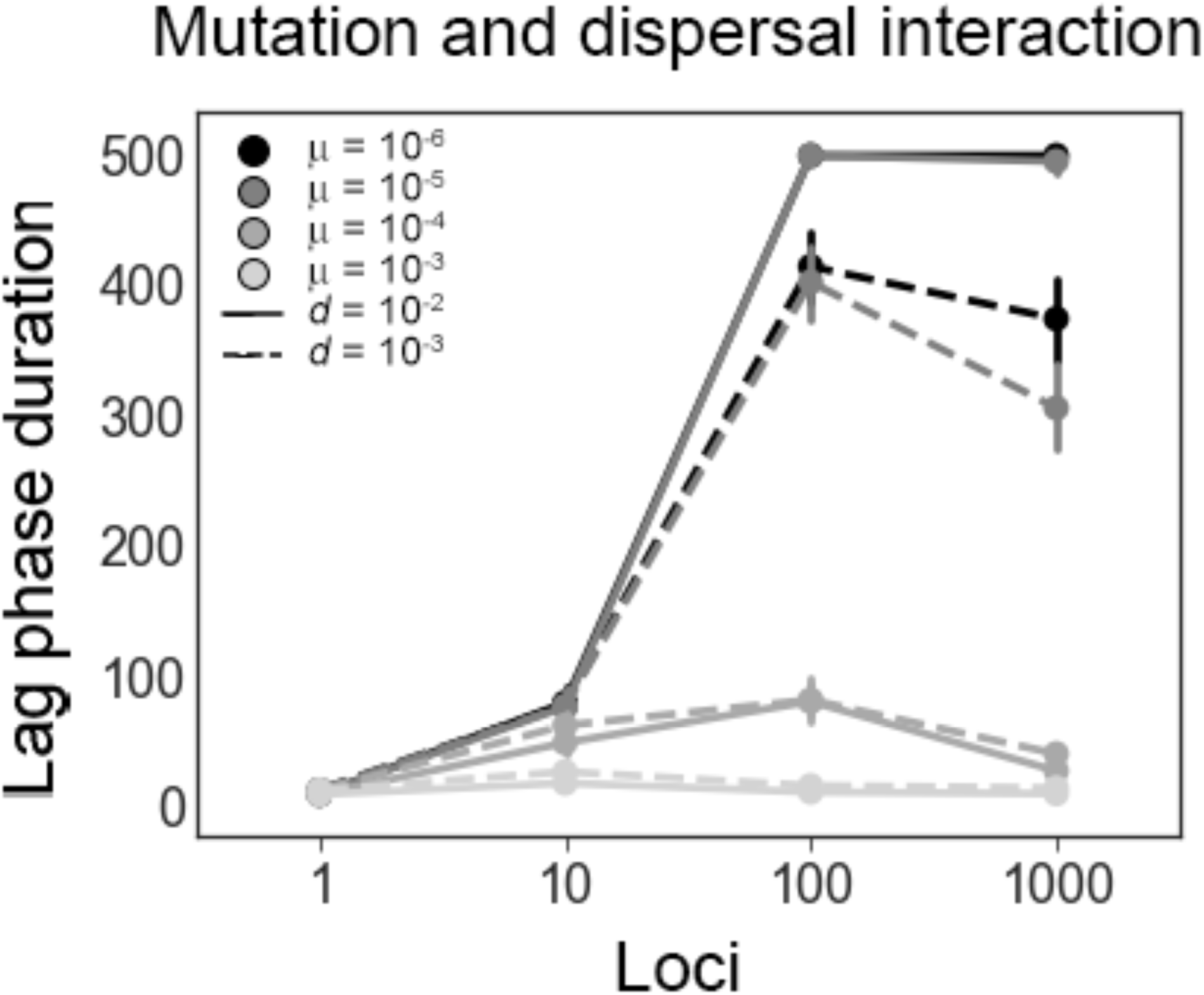
Mean colonization lag phase and 95% confidence intervals for each baseline architecture assuming different values of dispersal rate *d* (solid lines=10^−2^, dashed lines=10^−3^) and mutation rate μ (light grey=10^−3^, grey=10^−4^, dark grey=10^−5^, and black=10^−6^).

## Literature Cited

Abbott, R., Albach, D., Ansell, S., Arntzen, J.W., Baird, S.J.E., Bierne, N., Boughman, J., Brelsford, A., Buerkle, C.A., Buggs, R., et al. (2013). Hybridization and speciation. J. Evol. Biol. 26, 229–246.

Aikio, S., Duncan, R.P., and Hulme, P.E. (2010). Lag-phases in alien plant invasions: separating the facts from the artefacts. Oikos 119, 370–378.

Bock, D.G., Caseys, C., Cousens, R.D., Hahn, M.A., Heredia, S.M., Hübner, S., Turner, K.G., Whitney, K.D., and Rieseberg, L.H. (2015). What we still don’t know about invasion genetics. Mol. Ecol. 24, 2277–2297.

Comeault, A.A. (2018). The genomic and ecological context of hybridization affects the probability that symmetrical incompatibilities drive hybrid speciation. Ecol. Evol. 8, 2926–2937.

Comeron, J.M., and Kreitman, M. (2002). Population, Evolutionary and Genomic Consequences of Interference Selection. Genetics 161, 389–410.

Crooks, J.A. (2005). Lag times and exotic species: The ecology and management of biological invasions in slow-motion. Ecoscience 12, 316–329.

Crooks, J.A., and Soulé, M.E. (1999). Lag times in population explosions of invasive species: causes and implications. Invasive Species Biodivers. Manag. Based Pap. Present. NorwayUnited Nations UN Conf. Alien Species 2nd Trondheim Conf. Biodivers. Trondheim Nor. 1-5 July 1996 103–125.

Dlugosch, K.M., Anderson, S.R., Braasch, J., Cang, F.A., and Gillette, H.D. (2015). The devil is in the details: genetic variation in introduced populations and its contributions to invasion. Mol. Ecol. 24, 2095–2111.

Ellstrand, N.C., and Schierenbeck, K.A. (2000). Hybridization as a stimulus for the evolution of invasiveness in plants? Proc. Natl. Acad. Sci. 97, 7043–7050.

García-Ramos, G., and Kirkpatrick, M. (1997). Genetic Models of Adaptation and Gene Flow in Peripheral Populations. Evolution 51, 21–28.

Gilbert, K.J., and Whitlock, M.C. (2017). The genetics of adaptation to discrete heterogeneous environments: frequent mutation or large-effect alleles can allow range expansion. J. Evol. Biol. n/a-n/a.

Gilbert, K.J., Sharp, N.P., Angert, A.L., Conte, G.L., Draghi, J.A., Guillaume, F., Hargreaves, A.L., Matthey-Doret, R., and Whitlock, M.C. (2017). Local Adaptation Interacts with Expansion Load during Range Expansion: Maladaptation Reduces Expansion Load. Am. Nat. 189, 368–380.

Gomulkiewicz, R., Holt, R.D., and Barfield, M. (1999). The Effects of Density Dependence and Immigration on Local Adaptation and Niche Evolution in a Black-Hole Sink Environment. Theor. Popul. Biol. 55, 283–296.

Harrison, R.G., and Larson, E.L. (2014). Hybridization, Introgression, and the Nature of Species Boundaries. J. Hered. 105, 795–809.

Holt, R.D., Gomulkiewicz, R., and Barfield, M. (2003). The phenomenology of niche evolution via quantitative traits in a “black-hole” sink. Proc. R. Soc. B Biol. Sci. 270, 215–224.

Kirkpatrick, M., and Barton, N.H. (1997). Evolution of a Species’ Range. Am. Nat. 150, 1–23.

Marques, D.A., Meier, J.I., and Seehausen, O. (2019). A Combinatorial View on Speciation and Adaptive Radiation. Trends Ecol. Evol.

McVean, G.A., and Charlesworth, B. (2000). The effects of Hill-Robertson interference between weakly selected mutations on patterns of molecular evolution and variation. Genetics 155, 929–944.

Neuenschwander, S., Hospital, F., Guillaume, F., and Goudet, J. (2008). quantiNemo: an individual-based program to simulate quantitative traits with explicit genetic architecture in a dynamic metapopulation. Bioinformatics 24, 1552–1553.

Pfennig, K.S., Kelly, A.L., and Pierce, A.A. (2016). Hybridization as a facilitator of species range expansion. Proc. R. Soc. B Biol. Sci. 283, 20161329.

Reatini, B., and Vision, T.J. (2020) Data from: Genetic architecture influences when and how hybridization contributes to colonization. Dryad Digital Repository https://doi.org/10.5061/dryad.3xsj3txbz

Rieseberg, L.H., Archer, M.A., and Wayne, R.K. (1999). Transgressive segregation, adaptation and speciation. Heredity 83, 363–372.

Schierenbeck, K.A., and Ellstrand, N.C. (2008). Hybridization and the evolution of invasiveness in plants and other organisms. Biol. Invasions 11, 1093.

Seehausen, O. (2004). Hybridization and adaptive radiation. Trends Ecol. Evol. 19, 198–207.

Simon, A., Bierne, N., and Welch, J.J. (2018). Coadapted genomes and selection on hybrids: Fisher’s geometric model explains a variety of empirical patterns. Evol. Lett. 2, 472–498.

Tigano, A., and Friesen, V.L. (2016). Genomics of local adaptation with gene flow. Mol. Ecol. 25, 2144–2164.

Wangen, S.R., and Webster, C.R. (2006). Potential for multiple lag phases during biotic invasions: reconstructing an invasion of the exotic tree Acer platanoides. J. Appl. Ecol. 43, 258–268.

Yeaman, S. (2015). Local Adaptation by Alleles of Small Effect. Am. Nat. 186, S74–S89.

Yeaman, S., and Otto, S.P. (2011). Establishment and Maintenance of Adaptive Genetic Divergence Under Migration, Selection, and Drift. Evolution 65, 2123–2129.

Yeaman, S., and Whitlock, M.C. (2011). The Genetic Architecture of Adaptation Under Migration–Selection Balance. Evolution 65, 1897–1911.

